# Prediction and Design of Protease Enzyme Specificity Using a Structure-Aware Graph Convolutional Network

**DOI:** 10.1101/2023.02.16.528728

**Authors:** Changpeng Lu, Joseph H. Lubin, Vidur V. Sarma, Samuel Z. Stentz, Guanyang Wang, Sijian Wang, Sagar D. Khare

**Affiliations:** Institute for Quantitative Biomedicine, Rutgers - The State University of New Jersey, Piscataway, NJ; Department of Chemistry & Chemical Biology, Rutgers - The State University of New Jersey, Piscataway, NJ; Verily Life Sciences, Boulder, CO; Department of Statistics, Rutgers - The State University of New Jersey, Piscataway, NJ

## Abstract

Site-specific proteolysis by the enzymatic cleavage of small linear sequence motifs is a key post-translational modification involved in physiology and disease. The ability to robustly and rapidly predict protease substrate specificity would also enable targeted proteolytic cleavage – editing – of a target protein by designed proteases. Current methods for predicting protease specificity are limited to sequence pattern recognition in experimentally-derived cleavage data obtained for libraries of potential substrates and generated separately for each protease variant. We reasoned that a more semantically rich and robust model of protease specificity could be developed by incorporating the three-dimensional structure and energetics of molecular interactions between protease and substrates into machine learning workflows. We present Protein Graph Convolutional Network (PGCN), which develops a physically-grounded, structure-based molecular interaction graph representation that describes molecular topology and interaction energetics to predict enzyme specificity. We show that PGCN accurately predicts the specificity landscapes of several variants of two model proteases: the NS3/4 protease from the Hepatitis C virus (HCV) and the Tobacco Etch Virus (TEV) proteases. Node and edge ablation tests identified key graph elements for specificity prediction, some of which are consistent with known biochemical constraints for protease:substrate recognition. We used a pre-trained PGCN model to guide the design of TEV protease libraries for cleaving two non-canonical substrates, and found good agreement with experimental cleavage results. Importantly, the model can accurately assess designs featuring diversity at positions not present in the training data. The described methodology should enable the structure-based prediction of specificity landscapes of a wide variety of proteases and the construction of tailor-made protease editors for site-selectively and irreversibly modifying chosen target proteins.

## Introduction

Multi-specificity, the specific recognition and non-recognition of multiple substrates by protease enzymes, is critical for many biological processes and diseases^1–5^. For example, the selective recognition and cleavage of host and viral target sites by viral and host protease enzymes is critical for the lifecycle of many RNA viruses, including SARS-CoV-2^6–10^. Identifying proteolytic targets of proteases would, therefore, provide deeper insights into the mechanisms and biological functions of proteases^3,11^. As protease inhibitors are often designed to mimic substrates, the ability to predict substrates may also aid inhibitor design against novel viruses^12–15^. Furthermore, the ability to infer the global landscape of protease specificity, i.e., the set of all substrate sequence motifs that are recognized (and not recognized) by a given enzyme, would also enable the selection or design of bespoke proteases with specificities to degrade chosen biotechnologically-relevant or disease-related targets^16–20^.

Current experimental methods for protease substrate cleavage site identification involve assaying libraries of potential substrates for cleavage, one protease variant at a time^1,21–26^. Apart from being labor-intensive and time-consuming, only limited sampling of the protease:substrate sequence diversity is possible. Therefore, the development of rapid, cost-effective and generalizable computational approaches for precise prediction of specificity is valuable. Most current computational approaches for protease specificity prediction involve detecting and/or learning patterns in known substrate sequences using techniques ranging from inferred substitution matrices (e.g., PoPS^27^, GraBCas^28^, CAT3^29^, SitePrediction^30^, GPS-CCD^31^, PEPS^32^) to supervised machine learning (e.g., DeepCleave^33^, NeuroPred^34^, PROSPERous^35^, PROSPER^36^, iProt-Sub^37^, CASVM^38^, Cascleave^39^, and Pripper^40^). In some approaches (e.g., Procleave^41^), the accessibility of substrates depending on their solvent exposure and secondary structure assignment are also considered during prediction. We previously developed a supervised machine learning-based approach for specificity prediction in which protease-substrate interaction energy terms for the interface were considered^42,43^. We found that energetic terms play an important role in helping rank probabilities of cleavage. Similarly, inclusion of energetics in machine learning models was found to increase classification accuracy for identifying metalloenzymes^44^. While computational approaches have successfully guided experiments in finding novel cleavage sites and obtaining a better understanding of protease–substrate interactions, these black-box approaches do not provide physical/chemical insight into the underlying basis for a particular experimentally observed specificity profile, nor are they robust to substitutions in the protease, requiring re-training for every protease variant, thus making these unsuitable for guiding protease design. Thus, there is need for interpretable and generalizable computational models of protease specificity.

We reasoned that a more semantically rich model of specificity would encompass both substrate sequence and the explicit energetics of the protease-substrate complex. Specificity depends on the residue-level interactions between enzymes and substrates, and for this reason, we hypothesized that a high-resolution energetic representation of a protease-substrate complex will have a high predictive value. As the energies of a protein are a consequence of sequence, we anticipated that a sufficiently granular and accurate energetic representation may obviate the need for sequence features. Using energies rather than sequence-based models for protease specificity naturally enables the design of proteases by training on directed evolution trajectories aimed at altering protease specificity for benchmarking^45^. To encode the topology and energetic features of protease-substrate complexes for modeling specificity landscapes, here we develop a Protein Graph Convolutional Network (PGCN). PGCN uses experimentally derived data and a physically intuitive structure-based molecular interaction energy graph to pose specificity prediction as a classification problem. We find that PGCN consistently performs as well as or better than other previously-used machine learning models for substrate specificity prediction especially when using energy-only features. A single PGCN model can effectively predict specificities for multiple protease variants, and ablation tests enable identification of critical sub-graph patterns responsible for observed specificity patterns, highlighting the interpretability of the model. We then use PGCN to guide the design of protease libraries aimed at cleaving non-canonical substrates for TEV protease, and experimentally validate these predictions using a yeast surface display-based assay. Importantly, designs included residue positions and substitutions not present in the training set, speaking to the high generalizability of PGCN.

## Results

### Overview of PGCN

We present a protein graph convolutional network (PGCN), which models protein structures and their complexes as fully connected graphs encoding sequence and single-residue and pairwise interaction energies generated using Rosetta^46^. For the protease-substrate complexes, the substrate peptide is recognized by the protease for cleavage or rejection in the active site (**Figure 1A**). The enzyme-substrate graph (**Figure 1B**) is fed into a graph convolutional neural network, which outputs a probability of cleavage for a given complex (**Figure 1C**). Our protease specificity dataset consists of experimentally-determined cleavage information, i.e, lists of cleaved and uncleaved peptides for the wild type and variants of two viral proteases, NS3/4 protease of the Hepatitis C Virus (referred to as HCV protease in the following) (**Table S1A)**, and Tobacco Etch Virus (TEV protease) (**Table S1B**) obtained from Pethe et al.^42^ and Packer et al.^45^. The pools of experimentally confirmed cleaved and uncleaved substrates were randomly split into 80% training, 10% validation and 10% test datasets.

**Figure 1:**
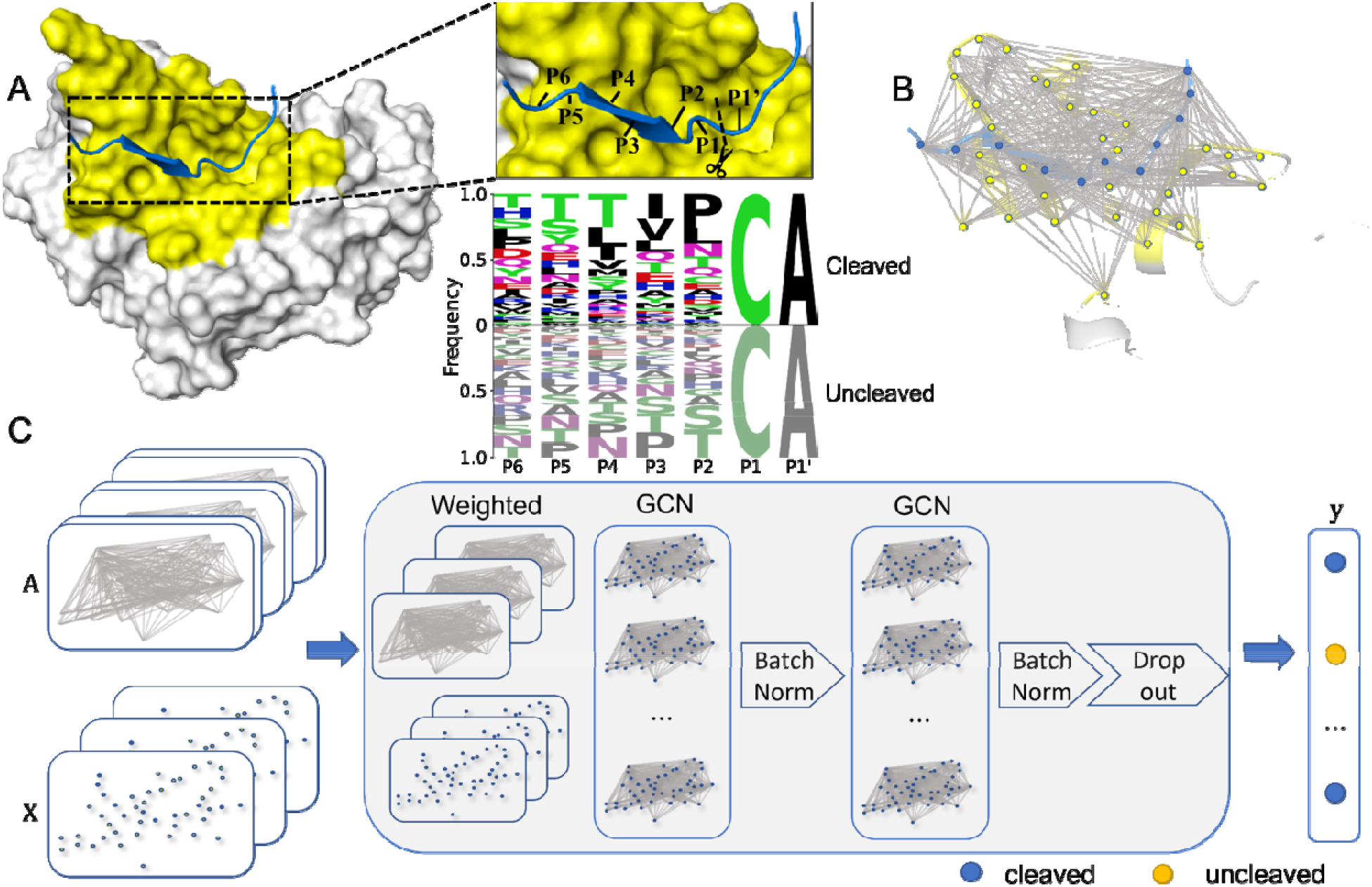
Architecture of PGCN A. Peptide substrate (blue) in the binding pocket (yellow) of HCV protease (grey). The 7-residue substrate spans P6 to P1’, with cleavage between P1 and P1’. The logo plot indicates the substrate sequences in the HCV training set, where P1 and P1’ were kept constant, and P6-P2 were variable. **B**. Molecular depiction of the nodes and edges as a graph. Each substrate (blue) and binding pocket (yellow) amino acid constitutes a node of the graph. Gray lines between pairs of residues denote edges between pairs of nodes. **C**. PGCN model architecture. Nodes are represented as a matrix of nodes and node features. Edges are represented as a tensor of node pairs and edge features, flattened by the weighted sum of overall edge features. The PGCN model ultimately outputs probabilities of the given substrate belonging to each class, cleaved and uncleaved.

### PGCN performs better than baseline ML models for HCV protease specificity prediction for various feature encodings

To evaluate the performance of PGCN predicting substrate specificity, we first trained and tested models for specificity landscapes of WT and three HCV variants, A171T, D183A, and R170K/A171T/D183A (**Table 1**). We further combined all HCV variant data and trained and tested a single PGCN model on this combined set to explore how sensitive PGCN is in discriminating specificity changes upon small structural changes in the protease.

**Table 1.** Summary of Input Dataset.

| Protease Variant | Training |  |  | Validation |  |  | Test |  |  | Total |
| --- | --- | --- | --- | --- | --- | --- | --- | --- | --- | --- |
|  | Cleaved | Uncleaved | Total | Cleaved | Uncleaved | Total | Cleaved | Uncleaved | Total |  |
| HCV WT | 1,566 | 4,307 | 5,873 | 175 | 559 | 734 | 191 | 544 | 735 | 7,342 |
| HCV A171T | 2,905 | 7,659 | 10,564 | 366 | 954 | 1,320 | 373 | 948 | 1,321 | 13,205 |
| HCV D183A | 3,538 | 5,953 | 9,491 | 422 | 764 | 1,186 | 390 | 797 | 1,187 | 11,864 |
| HCV Triple* | 2,496 | 2,974 | 5,470 | 315 | 369 | 684 | 324 | 360 | 684 | 6,838 |
| HCV | 10,404 | 20,995 | 31,399 | 1,319 | 2,606 | 3,925 | 1,338 | 2,587 | 3,925 | 39,249 |
| Combined |  |  |  |  |  |  |  |  |  |  |
| TEV<br>Combined | 2,111 | 2,229 | 4,340 | 259 | 283 | 542 | 238 | 305 | 543 | 5,425 |

\*Mutations made for HCV Triple: R170K, A171T, D183A

In benchmark tests, PGCN outperformed other ML models for all HCV variants (**Figure 2A**), achieving more than 90% test accuracy for all datasets, including the combined dataset. We evaluated PGCN performance using different metrics besides accuracy, including F1 score, Precision, Recall, Area under curve (AUC), and Average Precision (AP), all standard evaluation metrics for machine learning tasks with imbalanced data^47–50^. PGCN had the highest F1, Recall, and AP scores of the benchmarked methods (example: 93.53% F1, 90.44% Recall, 96.05% AP for A171T protease using sequence features only) (see details in **Table S2**).

**Figure 2:**
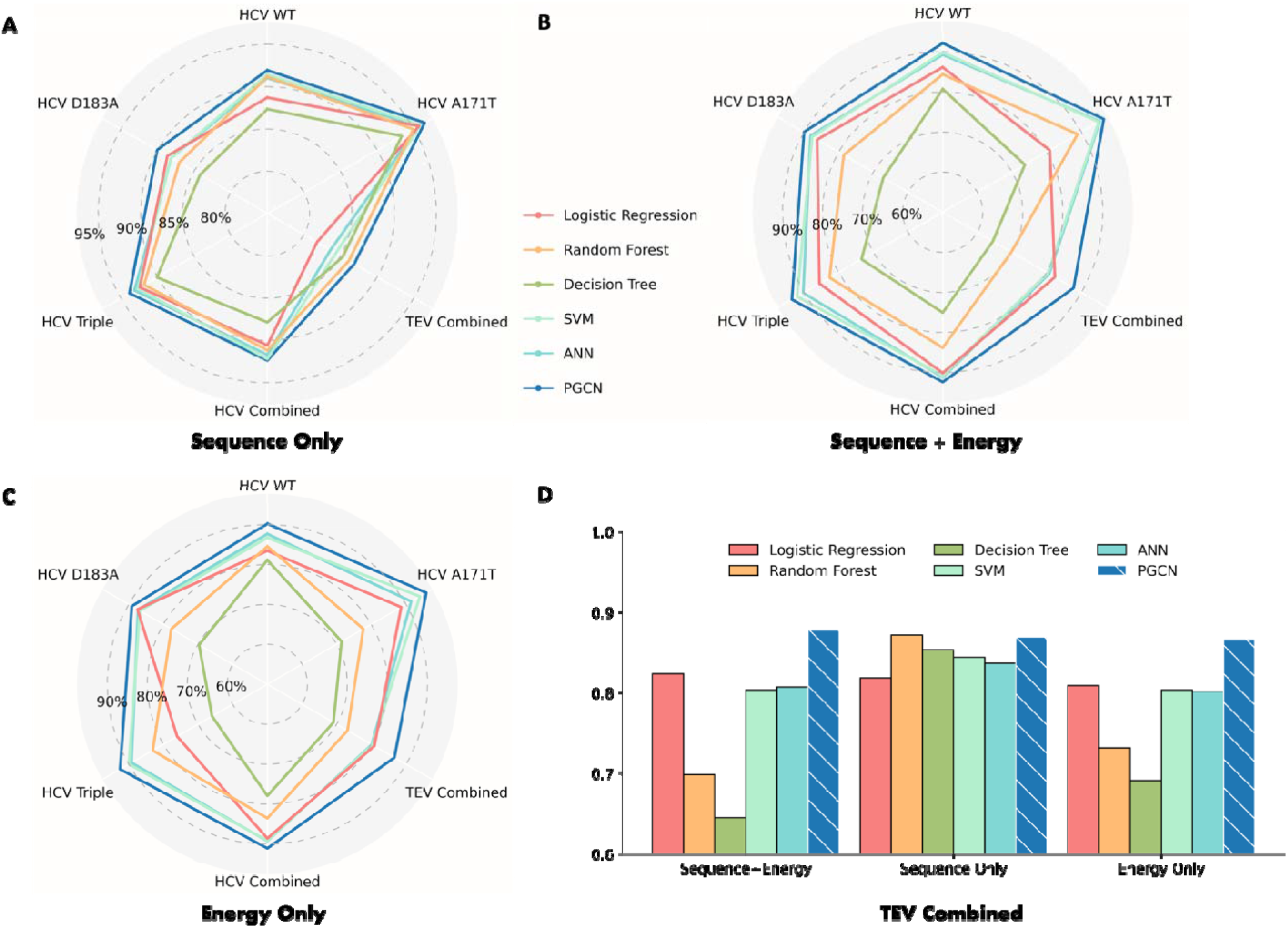
PGCN performance. We evaluate models on six datasets, four consisting of a single HCV variant (WT, A171T, D183A, or Triple (R170K, A171T, D183A)), with various substrates, and two, Combined, one of which pools the other four, and the other consists of 10 TEV variants. A-C) The radar plots show polygon patterns of average test accuracies across three seeds of benchmarked ML models (labeled in different colors) on the five datasets. The highest accuracy is on the polygon periphery. D) Accuracy barplot of models on TEV protease specificity data under different feature settings. Y-axis shows accuracies, truncated from 0.6.

We then evaluated PGCN’s performance when using energy features. In these tests, we used either Rosetta energy information only, or sequence and Rosetta energy information together as features used in PGCN. As shown in **Figures 2B and 2C**, PGCN always performed the best with either energy features only or complete sequence and energy features. This result is remarkable because previous energy-based scoring approaches for protease-peptide interactions, which involved weighted sums of different energy terms did not perform as well as sequence-based learning approaches^42,43^. A key difference between other energy-based models and PGCN is how calculated energies of interaction are used as features. In all models other than PGCN, energies are learned in simple linear combinations, while PGCN takes advantage of graph representation to encode intermolecular energies in an implicitly non-linear relationship. Therefore, our results show that graph-based convolution of individual energy terms is a promising approach for combining biophysical analysis and data-driven modeling in a way that addresses some of the limitations of each.

### PGCN performance in recapitulating TEV protease variant specificity with various feature encodings

Having demonstrated good performance in predicting substrate specificities when provided training data including a large pool of substrates and sparse protease diversity, we sought to evaluate PGCN performance in predictions involving greater protease diversity. Therefore, we trained PGCN on the engineered TEV protease dataset which had a larger set of protease variants than our HCV set, although fewer substrates per variant were experimentally assayed^45^ (**Table S3**). We trained on this TEV dataset using the same three sets of feature encodings, either sequence-only information, energy-only information, or complete sequence and energy information.

All ML models are able to learn some patterns for TEV data if considering sequence features only, but tree-based approaches, SVM, and ANN achieved lower accuracy when considering energy features (**Figure 2D**). As expected, PGCN’s performance is stable among different feature encodings, with accuracies of 86.86%, 86.62% and 87.72% when using sequence-only, energy-only, and sequence+energy features, respectively. PGGN takes advantage of encoding residue-level pairwise energies into edges of graphs that enable PGCN to learn the local environments of residues at each GCN layer. Furthermore, PGCN with complete sequence and energy features outperforms the models with reduced features (e.g., SVM), supporting our hypothesis that the prediction of protease specificity benefits from both sequences of substrates and physical energies of interaction between enzyme and substrate.

### Node-Edge Importance Analysis to Obtain Physical Insights from PGCN

One advantage of PGCN is that the nodes and edges correspond directly to physical amino acid residues and their relationships. Therefore, we reasoned that we could identify important residues and interactions by identifying nodes and edges found to be critical for PGCN performance. To identify the prediction strength of each graph component by PGCN, we perturbed feature values of each node (or edge) across all sample graphs and computed accuracy again (see *Node/Edge Importance in Methods*). The decrease in accuracy upon perturbation is used to measure the (relative) importance of node *i* (or edge *J*) in the PGCN graph.

We normalized the calculated importance of node/edge by the overall accuracy of the prediction and aggregated the normalized importance by feature type (node or edge) to see how the features used by PGCN for training affected the classification. There are two types of nodes (protease, substrate) and three types of edges (protease-protease, substrate-substrate and intermolecular) depending on the types of nodes that are connected by a given edge. When the sequence is the only feature (nothing on edges), as expected only peptide nodes contribute to accuracy for single-variant sets (**Figure 3A**). However, for datasets in which protease diversity is also sampled (“Combined” dataset in **Figure 3A**), protease nodes, typically sites of substitutions, are also detected as contributors to accuracy. When energy features are considered either solely or together with sequences, protease nodes make significantly greater contributions (**Figure 3A**), indicating that 1-body residue energies are highly sensitive to the changes in their environment. In the same vein, when the sequence information is excluded, the dependence on edge features increases while the overall accuracy of prediction is not significantly affected. Leveraging energy information allows broader attention to residue-residue interactions as more edges are deemed significantly important, as shown in **Figure S1**. These observations show that sequences are an abstraction that PGCN uses as a shortcut when available, but the same information can be learned from energy.

**Figure 3.**
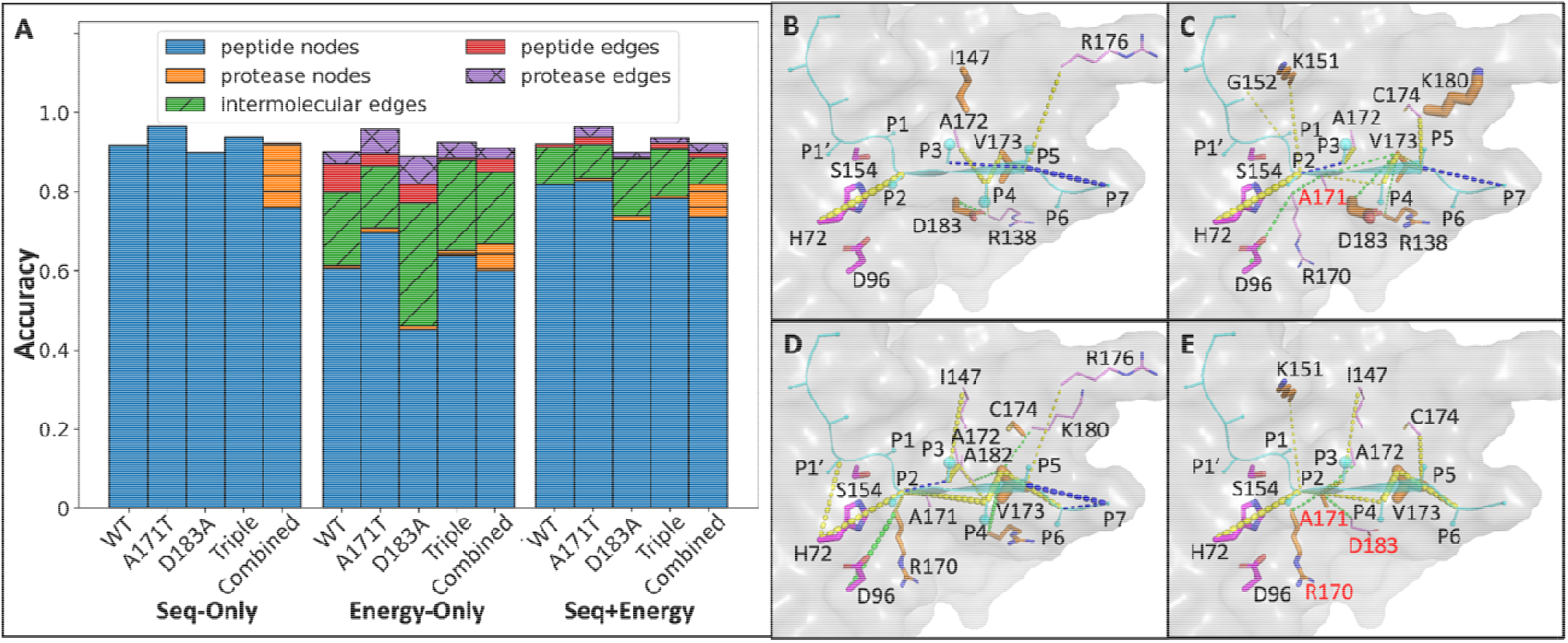
Node/edge importance analysis for HCV protease. A) Relative importance of major physical groups of nodes/edges. Node/edge importance is analyzed by the decrease in accuracy upon perturbation normalized by the original test accuracy of the PGCN model trained on each of five HCV protease variants using sequence-only S), energy-only (E), or both (S+E). All nodes and edges in the PGCN graph are grouped into five major physical groups, and each group’s total relative importance is aggregated as the sum of importance scores from nodes (or edges) of the specified type. B-E) Structural representations of node/edge importance for HCV protease (PDB ID 3M5N; 3M5L), calculated by the energy-only PGCN model trained on B) wild type; C) A171T; D) D183A; E) Triple data. Only nodes/edges whose relative importance scores are within the first 25% quantile are displayed in the zoom-in protease structure (grey), with the substrate (cyan) as the center and the catalytic triad (magenta) as the reference. Among those nodes that meet the criteria, the relative importance levels of protease nodes (orange) are shown by the thickness of corresponding residue side chains, while that of peptide nodes (cyan) is reflected by the sizes of corresponding residue spheres at CB. For those important edges that meet the criteria, different groups are highlighted in different colors, including peptide edges (blue), protease edges (green), and intermolecular edges (yellow). All residues related to node/edge importance have labels in residue identifiers with one-letter codes (colored in red if mutation sites), including protease residues that are only related to edges (violet).

Next, we visualized the positions of important nodes and edges in the HCV protease structure. For the wild type and each variant protease, a key edge was found to be between the P2 residue of the substrate and the catalytic base H172, which presumably reflects the proper positioning of the substrate in the active site. We also observed that some important nodes/edges were different between wild-type and variant proteases. For example, the protease edge R138-D183 is prominent for the wild type (**Figure 3B**), but it is not a significant interaction when either A171T (**Figure 3C**) or D183A (**Figure 3D**) or the Triple variant A171T, D183A and R170K (**Figure 3E**). When D183A mutation is introduced (**Figures 3D-3E**), side chain orientations of some intermolecular edges, e.g., P6-R138, were different even though the protease node D183A itself did not significantly influence classification for substrate specificity. Conversely, we found that some other intermolecular edges such as P3-I147, P2-A171(T), P4-A171(T), P4-V173, P6-V173 are at least two times more important for models trained on D183A and Triple variants than those for models trained on the wild type and A171T data (**Table S4**). Also, protease node R170(K) shows its importance only if D183A is mutated (**Figures 3D-3E**). However, the important edge and node lists are not additive: for example, edges 138-173 and 96-170 are insignificant for the Triple variant (**Figure 3E**), while they are two of the most important protease edges for the model trained on D183A (**Figure 3D**). When taking the model trained on the HCV combined set into consideration, although most of the important nodes and edges for single variant sets are equally important for the combined set (such as node V173, edge P2-H72, P4-V173, etc.), there are also some important nodes/edges that are only useful for the prediction of substrate specificity within individual variants or the wild type, such as P3-I147 (**Figures S1H-S1I**). Thus, the model is able to classify to approximately the same level of accuracy using overlapping but distinct sets of node and edge features.

Similar trends were apparent in the node/edge sensitivity analyses for TEV protease predictions (**Figure 4A**). Several positions that are the sites of amino acid substitutions in the TEV variants had high importance, such as D148, S170, and N177 (**Figure 4B**). Strong signals of some important interactions were also identified. For example, the interaction between the P2 residue of the substrate and the catalytic base H46 is consistently important across all TEV variants and the wild type (**Figure 4C**), the same as in the HCV prediction described above. In addition, several other edges, e.g., P3-S170 (**Figure 4C**), intra-protein edges around the S3 pocket (**Figure 4D**) and the S1 pocket (**Figure 4E**) were also found to be of high importance.

**Figure 4.**
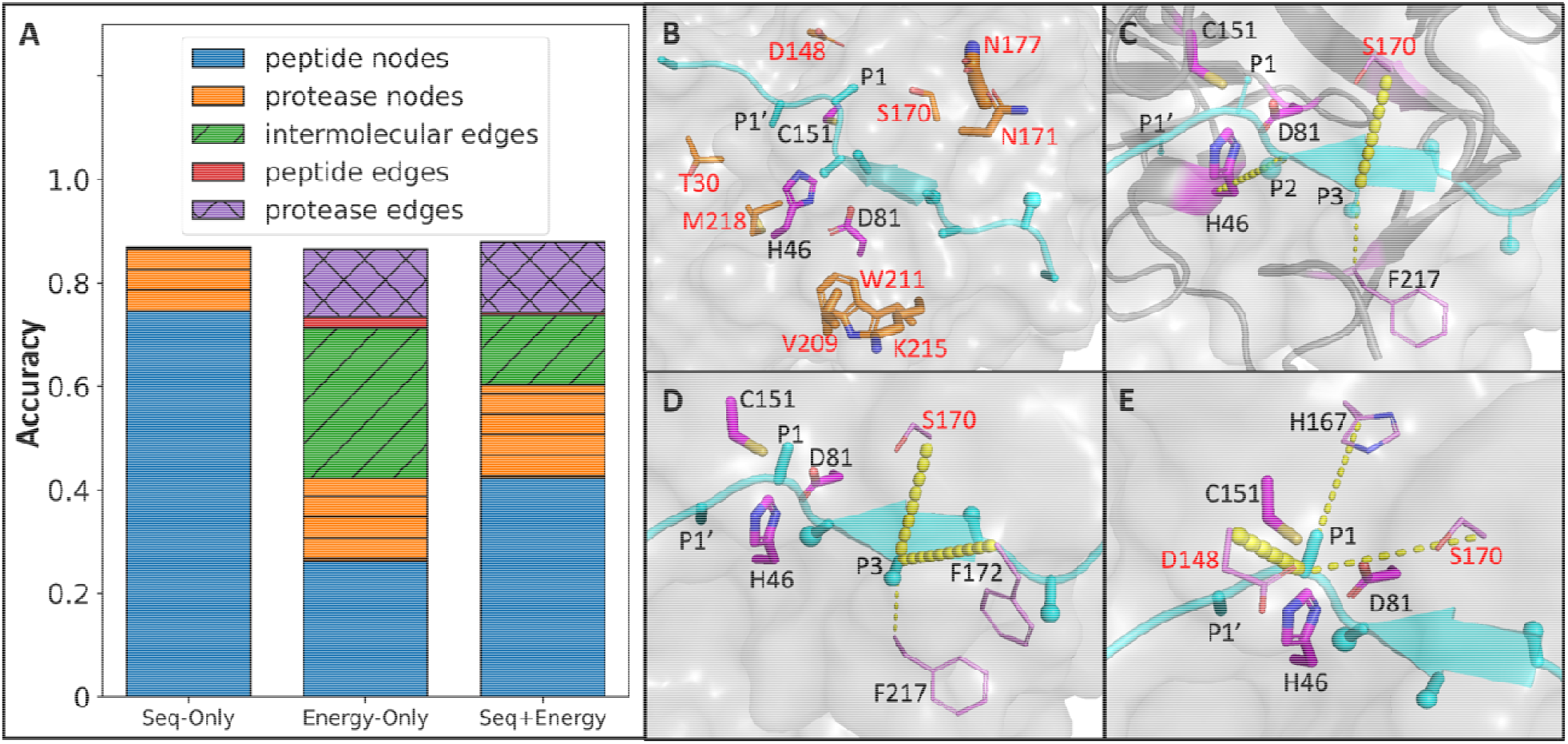
Node and edge importance contribution for TEV. A) Relative importance of major physical groups of nodes/edges. B-E) Partial structural representations of node and edge importance in the context of TEV (PDB ID 1LVB; 1LVM) to show: B) important protease nodes; C) intermolecular edges that presumably form hydrogen bonds; D) the subpocket surrounding P3, including P3-S170, P3-F172, P3-F217; E) the subpocket surrounding P1, including intermolecular edges P1-D148, P1-H167, P1-S170. The same color setting is used as in **Figure 3**. The catalytic triad in each structure is just the reference of relative position. All mutations are highlighted in red.

Taken together, the analyses for HCV and TEV protease variants follow a series of general rules. First, the intermolecular edge between P2 and the histidine in the catalytic triad (P2-H72 for HCV in **Figures 3B-3E**, and P2-H76 for TEV in **Figure 4C**) is always one of the most important edges, reflecting the proper positioning of the substrate in the active site. Second, those interactions that presumably provide the H-bonds that template the substrate into the required β-sheet conformation were also identified as prominent edges, such as P4-V173 for HCV (**Figures 3B-3E**), P3-S170, P4-F217 for TEV (**Figure 4C**). This is consistent with the well-known observation that protease substrates always adopt an extended conformation in the active site with beta-sheet complementation^51^. Some important nodes/edges form interconnected clusters that are consistent with the canonical substrate binding pockets (S pockets), for example, the S3 subpocket for TEV includes residues 170, 172, 217 (**Figure 4D**); the S1 pocket for TEV including interactions with 148, 167, 170 (**Figure 4E**). Thus, we argue that the PGCN models have learned to discriminate between cleaved and uncleaved substrates based on criteria that can have an interpretable biophysical basis in some cases, and reflect non-obvious statistical relationships between various interactions in other cases.

### Experimental Evaluation of PGCN generalizability using protease design

To test if PGCN is able to generalize its classification ability to protease variants that are not in the training dataset, we turned to TEV protease specificity design. Wild type TEV protease demonstrates a preference for an ENLYFQ/X motif at the P6-P1’ positions of its canonical substrate (X = A,G,S)^45,52,53^. We aimed to design proteases against altered substrates with single residue substitutions within the canonical recognition motif (P6: KNLYFQ/A, P2: ENLYYQ/A). A substitution of K at P6 or Y at P2 resulted in no cleavage of the substrate by wild-type TEV (**Figure S2**), providing a well-posed problem for designs using PGCN – given a set of designs, predict which designed variants would lead to cleavage of P6 and P2 target substrates.

We applied Rosetta-based computational design (*Computational design process for TEV protease in Supplementary Material*) to propose sequences (4,320 P6-targeted designs and 280 P2-targeted designs) that included stabilizing interactions with the target substrates (**Figure 5A**). We then used our pre-trained TEV protease model to score designs, and identified those with a high predicted probability of cleavage (**Figure 5B**). PGCN selected 200 of 280 P2-targeted designs and 126 of 4,320 P6-targeted designs with high predicted cleavage probability. Based on visual inspection, we selected 96 energetically favorable designs targeting P6 and 18 energetically favorable designs targeting P2 for further evaluation. To cost-effectively test these designs in experiments as well as generate negative (uncleaved) examples for classification, we identified amino acid substitutions that were enriched in the selected designs (**Figure S3**) and generated a combinatorial library sampling these protease substitutions. The library of selected designs was subjected to our previously reported version of the YESS (Yeast endosomal sequestration screening) assay^54^ (**Figure 5C**). Briefly, the assay detects cleavage by the presence/absence of HA and FLAG tags on either side of the chosen substrate using anti-tag fluorescently-labeled antibodies, and FACS is used to isolate pools of “cleaved” and “uncleaved” cells. Several individual colonies from plating the cleaved and uncleaved pools of the P6- and P2-targeted TEV libraries were clonally validated (**Figure 5D**). We selected 19 designs for clonal validation, 9 from the cleaved pool and 10 from the uncleaved pool. As shown in **Figure 5E**, all 9 cleaved designs are correctly predicted by PGCN with high confidence (probability > 0.75) whereas 7 out of the 10 uncleaved designs are correctly predicted. Predictions are robust across 10 different Rosetta FastRelax-generated models of designs (**Figure S4A**).

**Figure 5.**
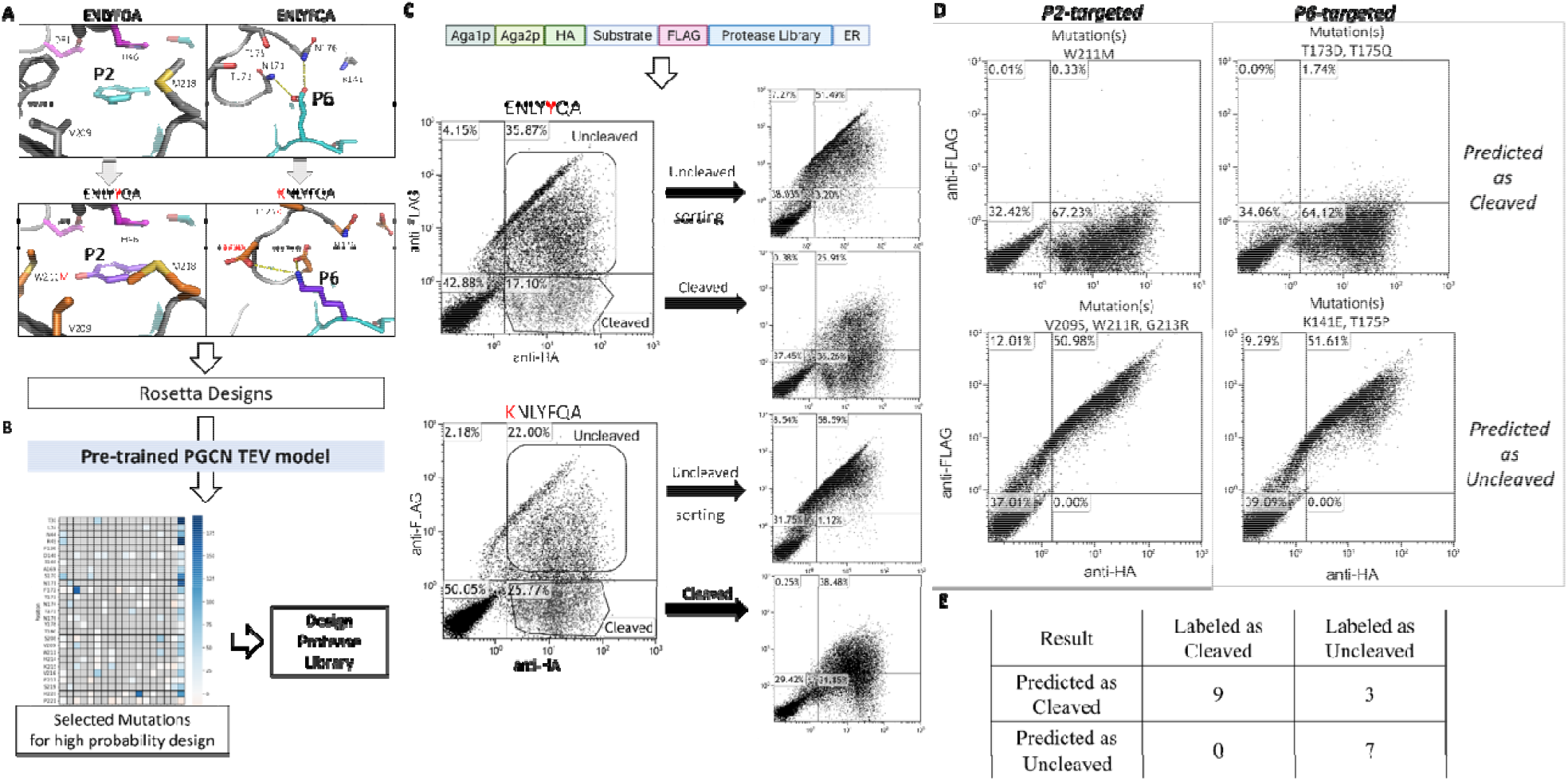
Pipeline for TEV protease design, including procedures of A) computational design, B) PGCN prediction, and C) yeast-based assay testing using FACS, and D) flow cytometry-based analysis of individual colonies. A) S2 and S6 Pockets corresponding to altered substrate residues were inferred from the crystal structure of a TEV protease-substrate complex, and redesigned using Rosetta. B) Rosetta-generated designs were evaluated using a pre-trained PGCN model and mutations enriched in high-scoring designs were identified and used to generate combinatorial protease libraries. C) These libraries were screened using the YESS assay with P6 and P2 variant substrates, and pools of cells corresponding to cleaved and uncleaved protease:substrate variant pairs were isolated using FACS. D) Individual colonies isolated from cleaved and uncleaved pools were tested using flow cytometry. 2-D scatter plots for positive P2-targeted (top-left) and P6-targeted (top-right) designs and negative designs (P2: bottom-left, P6: bottom-right) are shown. E) Comparison table of PGCN prediction and experimental results on 19 clonally-tested designs.

## Discussion

Current computational methods for protease specificity prediction largely rely on statistical pattern detection in datasets of known substrates and non-substrates for learning specificity, and are therefore not generalizable for use in protease design, and do not provide insights into the underlying biophysical bases of substrate-protease molecular recognition. We developed PGCN to include residue-level energies as features and decompose the Rosetta-computed energies into the node and edge features. PGCN shows high accuracy in classification tasks for HCV and TEV protease variants. A PGCN model utilizing sequence and energy-based features and trained on experimental data for TEV protease was used to evaluate and select designed TEV protease variants for cleaving non-canonical. Evaluated protease diversity included residue positions and/or amino acid substitutions not present in the training dataset. Experimental validation showed that PGCN scoring led to high accuracy selection of cleaved designs. Thus, PGCN was able to be generalized and our studies show proof-of-principle for the challenging task of protease design using ML-enabled structure-aware computational modeling.

The physically-based graphical structure of PGCN enables overcoming to some extent the black-box nature of ML methods as applied to protein modeling. Sensitivity analysis of nodes/edges that are mapped to a residue (a node) or the link between two residues (an edge) helps identify residues and pairwise residues that are most influential for selectivity, suggesting that the latent space of the PGCN model is rich and has the potential to further our understanding of molecular bases of protease selectivity. For example, identified important intermolecular edges influence the relative placement of the scissile bond with respect to the catalytic base, a key geometric requirement for the acylenzyme formation step protease hydrolysis. Furthermore, PGCN identified sub-graphs or networks of interacting residues that are key for specificity in recognizing a single peptide residue. Current implementation of PGCN gives as output probabilities for a binary classification, thus the results are qualitative. A quantitative or semi-quantitative prediction (i.e., ranking with respect to a known variant) of the catalytic parameters of the enzyme may become possible with protease activity datasets of sufficiently large size generated using experiments in which proteolysis is measured as a function of time. Due to its physical grounding, PGCN may be generally applicable in the specificity prediction and design of other proteins that bind peptide substrates, especially when only small experimental datasets (a few hundred or thousand peptide substrates) are available^55^. As PGCN only needs the structure of one protein:peptide complex as input, the availability of predicted structures of protein-peptide complexes opens the door to tackling specificity modeling on a large scale, provided accurate complex structures can be built^56,57^. These efforts are currently ongoing in our laboratory.

## Materials and Methods

### Protease Specificity Data

#### HCV protease

The experimental dataset for HCV protease was obtained in previous yeast surface display experiments conducted in our lab using yeast surface display coupled with deep sequencing^42^. This method allowed for rapid sampling of many candidate P6-P2 sequences with a given protease (WT HCV or one of three variants shown in **Figure S5A**) and determining each of those substrates to be either cleaved or not cleaved. Specifically, it sampled 7,342 substrates for wild type (1,932 of which were confirmed as cleaved), 13,208 substrates for A171T variant (3,644 of which were confirmed as cleaved), 11,864 substrates for D183A variant (4,350 of which were confirmed as cleaved) and 6,838 substrates for R170K+A171T+D183A variant (3,135 of which were confirmed as cleaved). All data points for HCV proteases are combined into a new single dataset (named *HCV Combined*). For each protease variant, substrate identity within each pool is less than 80% to avoid overfitting to the input because of data redundancy with a number of samples for each variant shown in **Table 1**. Experiments enable the sampling of P6-P2 substrates across all amino acids (**Figures S5B-S5F**). As the datasets for WT, A171T, and D183A are imbalanced (the number of uncleaved samples is at least twice of the cleaved samples), metrics besides accuracy, such as precision, recall, F1 score, etc., were also considered during the analysis.

#### TEV protease

The experimental dataset for TEV protease was obtained from directed evolution and deep sequencing profiling data of the phage substrate display collected in Packer et al, 2017^45^. Briefly, substrates of nine designed TEV variants (**Table S3**) were profiled based on single-mutation substrate libraries, and each variant has between 4,000 to 6,000 substrates, including 2,000-3,000 cleaved sequences. L2F variant and wild type (WT) variant were also profiled based on a triple-mutation substrate library (three substrate positions are randomized simultaneously). Up to ∼55,000 cleaved sequences and ∼80,000 uncleaved sequences for the L2F variant were obtained, while ∼30,000 cleaved sequences and ∼40,000 uncleaved sequences were obtained for the WT. However, due to the noise in the high-throughput sequencing assays, we observed considerable overlap between the cleaved and uncleaved pools for all variants. Therefore, we developed a data processing pipeline (**Figure S6**) to preprocess raw deep sequencing data to identify non-overlapping substrate sets (*TEV Deep Sequencing Data Processing in the Supplementary Methods)*. Sequence data were filtered based on empirical threshold to minimize overlap between cleaved and uncleaved populations (**Figures S7-S10**), resulting in a final tally of 5,425 high-confidence substrates among ten TEV protease types (including the wild type or variants), 48% of which are determined to be cleaved.

Observed substitutions in the engineered TEV variants are dispersed within the TEV protease structure (**Figure 6A**). Additionally, there is variation in both P1 and P1’ positions of substrates shown in **Figure 6B**, in contrast with HCV data. Other than Q/S, other P1/P1’ combinations, such as H/I, N/S, H/G, and Q/W, have been included. The top frequent amino acids at all positions in the cleaved population are the same as in the uncleaved population, and those from P6 to P1’ consist of ENLYFQ/S, known as the canonical sequence of TEV protease. Thus, the differences between cleaved and uncleaved sets are subtle.

**Figure 6.**
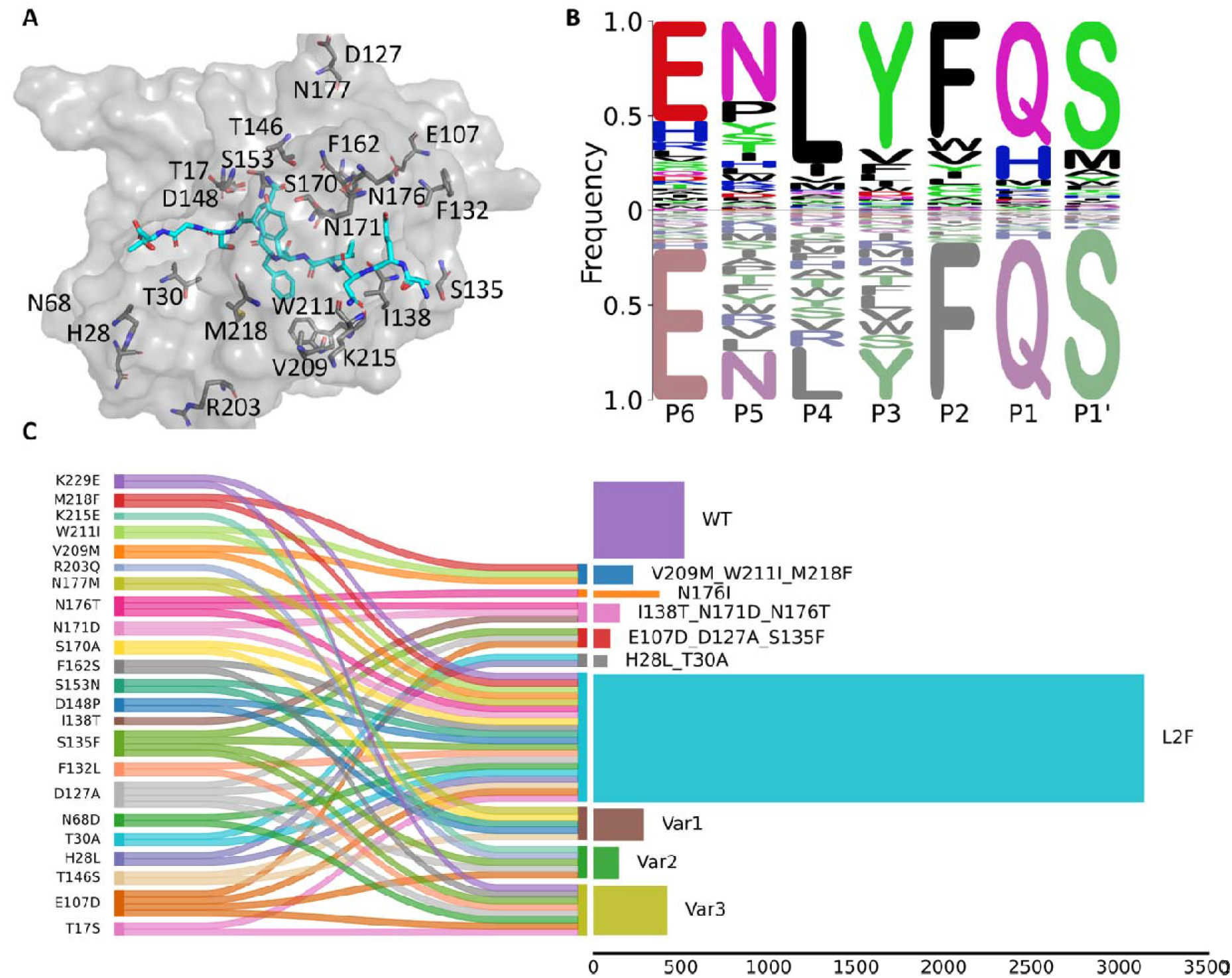
TEV input data variety. A) TEV protease mutation sites (grey) are shown all together in the TEV wild-type protease structure, located around the substrate (cyan). B) Substrate sequence logos for TEV input data from P6-P1’ positions. X-axis splits data into cleaved (above x-axis) and uncleaved (below x-axis) populations, where the higher frequency of amino acids appeared in the cleaved population, the higher it is located in the logo plot; the lower if considering the uncleaved population. C) Sankey diagram of mutation sites for TEV variants, combined with a horizontal barplot, showing the number of samples for different TEV variants (along x axis). Variants are named based on their mutation sites except for the following four variants: Var1 (T146S_D148P_S153N_S170A_N177M), Var2 (E107D_D127A_S135F_R203Q_K215E), Var3 (T17S_N68D_E107D_D127A_F132L_S135F_F162S_K229E), and L2F. See **Table S5** for complete mutation sites in the table.

In terms of the degree of protease variation in the TEV dataset, up to 23 different substitutions are present in comparison with the TEV wild-type protease (**Figure 6C, left**). As depicted in the Sankey diagram (**Figure 6C)**, among 10 TEV protease variants (including wild type), R203Q, K215E, and I138T are present in one variant; T17S, H28L, T30A, N68D, F132L, T146S, D148P, S153N, F162S, S170A, N171D, N177M, V209M, W211I, M218F (cyan) and K229E are present in two variants; N176T (orange) is present in three variants; E107D, D127A, S135F (magenta) are present in four variants. L2F variant has the largest number of amino acid substitution sites among all variants and accounts for the majority of the dataset.

### Protease-Substrate Complex Model Generation Using Rosetta

For HCV protease, models of protease-substrate complexes were based on crystal structures of inhibitor-bound (PDB code: **3M5N**) and peptide-bound inactive (PDB code: **3M5L**) variants of HCV protease^58^. The structures were superimposed, and the peptide substrate of 3m5l was copied into the unmutated active site of 3m5n, replacing the inhibitor, and then the complex was minimized using the Rosetta FastRelax protocol^59^ with coordinate constraints on all C□ atoms. A similar process was used with crystal structures of inhibitor-bound (PDB code: **1LVM**) and peptide-bound inactive (PDB code: **1LVB**) for TEV^52^.

Mutant complexes were generated by substituting the appropriate residues in the wild-type (WT) complex and then minimizing all interfacial side chains (interfacial residues were defined as those with Cα-Cα distance < 5.5Å, or with distance <9Å and Cα-Cβ vectors at an angle <75°) with an immobile backbone.

We generated models for each possible substrate (3.2 × 10^5^ possibilities within P5-P2) as part of a complex including the peptide from P7 to P4’ bound to each of the four HCV variants and the peptide from P7 to P3’ bound to each of the ten TEV variants in PyRosetta^23^, using the Rosetta FastRelax protocol^59^. Coordinate constraints minimized the movement of C α for the peptide backbone, and the protease backbone was held fixed. Distance, angle, and dihedral constraints were applied to the catalytic triad (H72, D96, and S154 for HCV and H46, D81, and C151 for TEV) and the P1 and P1’ residues to enforce the catalytic geometry required for cleavage. A single FastRelax trajectory was performed for each protease-substrate complex.

### Protein Graph Representation

The protein complex obtained from Rosetta modeling was encoded as a fully connected graph (i.e., there is an edge between every pair of nodes). The nodes of this graph are the amino acids of the substrate and the binding pocket of the protease, and the edges represent the pairwise residue interactions between nodes (**Figure 1B**). Each node contains the features of a single residue, including a 1-hot encoder for amino acid type, a binary term to indicate whether a residue is part of the substrate or the protease, and all 1-body Rosetta energy terms^60^. Each edge contains relational features between a pair of residues, including binary indications of whether the edge is between a substrate and a protease residue and whether the residues are covalently bonded (i.e., adjacent residues in sequence), and all Rosetta 2-body energy terms. Node and edge features are listed in **Table S6**. Since residues that are far away from the substrate are less likely to influence specificity and more likely to introduce noise into the model, we only consider a sub-graph including the substrate and interfacial residues that are within 10 Angstroms from the substrate (see **Table S7A/S7B** for node indices for HCV/TEV input graphs), selected based on proximity and sidechain orientation toward the substrate.

The node features for HCV protease are encoded into a *N* × *F* node feature matrix *X*, where *N =* 34 is the number of nodes, and *F=* 20 + 8 is the number of node features if combining both sequence and energy features for HCV data. The edge features are encoded as a *M*-length list of *N* × *N* edge feature matrices, where *M=* 8 is the number of edge features.

For TEV data, the number of nodes is *N=* 47, while node and edge feature encodings are the same as HCV data. To better introduce the architecture of our model, we use HCV data under combined (Sequence + Energy) feature encoding as the reference in the following paragraphs.

### Transformation of Energy Features

Rosetta energies are negative for favorable interactions and positive for unfavorable interactions and penalties. However, for learning, it is preferable if all features are positive values when updating node weights during convolution. To accomplish this conversion, we performed a two-step transformation. First, we transformed each element *e*_*i,j,q*_ in the edge tensor to a modified Boltzmann weight, 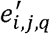 (**Equation 1**), where *i* and *j* are the nodes comprising the edge and *q* is the edge feature, numbered 1 to 8.

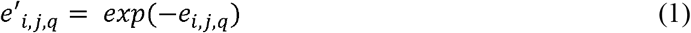

This converts negative Rosetta energies to larger positive values and positive Rosetta energies to small positive values, thereby assigning more appropriate weights to stabilize interactions. Second, since each edge of the graph is shared by two nodes, we followed Kipf^61^ to further normalize each edge weight by degrees of two ends of nodes to match normalized Laplacian. The normalized Laplacian is written in the matrix format, which is similar to **Equation 2**,

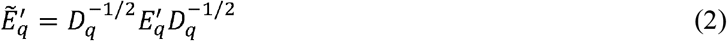

Herein, 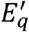 represents *q*th edge feature matrix, and each element of the matrix 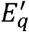 is 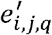, derived from **Equation 1**. *D*_*q*_ is the diagonal degree matrix of 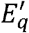, of which diagonal element is the sum of *q* th edge features for edges that are linked to a specific node.

After the transformation and normalization, the original edge data is transformed into a list of M matrices labeled by 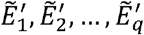. Each matrix is of the size *N* by *N*, where *N=* 34 denotes the number of nodes. Next, PGCN transforms the *M=* 8 edge matrices to a weighted sum *N* × *N* matrix *E*. In other words, 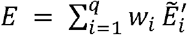 where *w*_1_, *w*_2_,..,*w*_*q*_ are learned weights. Those weights are initialized within the range of [0,1) and updated throughout the training process.

### Protein Graph Convolutional Network (PGCN)

After PGCN has transformed edge feature matrices into the matrix *E* with the size of *N* by *N*, each node from the node feature matrix *X* with the size of *N* by *N* is able to learn from all other nodes based on weights assigned by edge features. E matrix is used as the weight matrix for convolution. Then the weight matrix *E* is fed to a GCN^61^ layer (**Figure 1C**), which in fact results in that each edge feature matrix 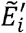 performs matrix multiplication with node feature matrix *X*, and independent multiplication outputs are concatenated and linearly transformed to *N* by *F* dimension, which aligns with the idea of multi-head attention^62^.

To reduce computational complexity, we used two GCN layers, and the number of hidden nodes of the second layer is the same as the first GCN layer. Therefore, the output of two GCN layers is

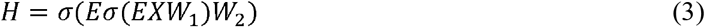

where *σ* is the nonlinear activation function^63^, here we use *σ = ReLU(x) = max(x*, 0) element-wise applied on each output of the GCN layer, *W*_1_, *W*_2_ are learned weight matrices both with the size of *F* × *C* for hidden layers with *C =* 20 feature maps. Each GCN layer is followed by a BatchNorm layer^64^, which aims at avoiding slow convergence.

Next, PGCN drops out a proportion of hidden nodes over nodes to avoid the overfitting problem^65^. Finally, PGCN flattens the output matrix *H* ∈ *R*^*N*×*C*^ from the dropout layer into a one-dimensional vector *H’* ∈ *R*^1×*NH*^. Then we transform *H’* to the expected dimension by applying a linear layer,

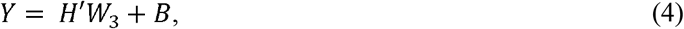

where *Y* ∈ *R*^1×1^ is the output, *W*_3_ ∈ *R*^*NH*×1^ is the learned vector, *B* ∈ *R*^1×1^ is the learned bias. Herein, we could apply the sigmoid as the activation function to calculate probabilities of being cleaved/uncleaved.

We followed a previously described approach to initialize weights^25^. For training, PGCN does backpropagation to update all parameters mentioned above and the set of tuning hyperparameters: batch size, learning rate, dropout rate, and weight decay coefficient. The weight decay coefficient is a part of the L2 regularization term that multiplies the sum of learned weights for the anti-overfitting procedure. Learned parameters are updated through epochs. The trained PGCN model is used for testing, in which test data pass through each layer of the PGCN model but skip the dropout process. The loss function is cross-entropy loss. PGCN is trained on training datasets using PyTorch, and tested on validation sets for hyperparameter tuning. PGCN performances are reported on test datasets.

### Comparison with baseline Machine Learning methods

We compared the prediction performance obtained for the PGCN with that obtained from five other machine learning (ML) methods. We used the Scikit-learn 0.20.1^66^ to implement logistic regression (lg), random forest (rf), decision tree (dt), support vector machine (SVM) classification, and Tensorflow 1.13.1^67^ for artificial neural network (ANN). The ANN model in this experiment is a one-layer fully connected neural network with 1,024 hidden nodes and allows a dropout rate between 0.1-0.9. To better compare performances between PGCN and other ML models, energy features are formed by residue-level energies, including single residue energies and pairwise energies flattened into a 1-dimensional vector, together with the protease type identifier (encoded in 10-dimensional one hot encoder) and sequence one hot encoder. In this case, PGCN and other ML models have the same feature encoding on energies to ensure proper comparison. We also followed the same rule as PGCN of splitting data into training, validation, and test datasets.

### Node/Edge Importance

We would like to derive biological insights about important residues or relationships between pairs of residues that contribute to the discrimination of cleaved and uncleaved substrates in PGCN. Since the test graphs for a PGCN model all come from the same protein family, they share the same graph structure. Therefore, we can discover the important residue (or a pair of residues) of the same node (or edge) across all test graphs. To efficiently determine the importance of a specific node (or edge), we perturbed values of each node feature for the same node (or edge feature for the same edge) across all test samples and inspected how much the test accuracy drops. By doing this, we avoid re-training the PGCN and the time complexity of perturbation is *0*(1). By following the procedure above, we were able to evaluate the importance of all nodes and edges on test graphs. We further normalized the change of accuracy by (Original_Accuracy – Perturbed_Accuracy)/Original_Accuracy.

### Combinatorial library preparation and yeast-surface display

Oligonucleotides containing degenerate codons at positions K141, N171, T173, T175, N176 (P6-targeted residues) or V209, W211 and M218 (P2-targeted residues) of the TEV protease sequence were purchased from IDT Inc. Application of degenerate codons increased the theoretical library size from 96 to 432 in the P6 targeted library and from 18 to 48 in the P2 targeted library. The double-stranded insert DNA sequence (varying from 150-300 bp in length) coding for the combinatorial amino acid library was assembled through overlap assembly PCR followed by agarose gel extraction and column purification. The integrity of the assembled insert was verified by Sanger sequencing through Genewiz Inc.

An LY104 vector backbone (obtained from Y. Li, B. Iverson, and G. Georgiou at University of Texas at Austin) containing the gene sequence for TEV protease and P6-P2 region of the corresponding substrate was linearized through PCR with primers to create sufficient overlap with the insert sequence for effective homologous recombination. Electrocompetent EBY100 yeast cells were transformed with the DNA library through electroporation on a Micropulser(tm) electroporation apparatus at 1.8kV. The cultures were grown overnight in selective dextrose casamino acid (SD-CAA) media. While the O.D of the cultures was less than 6, the cells were resuspended in selective galactose casamino acid induction media to induce display of the constructs on the surface of yeast.

### Library FACS and cytometric analysis of individual designs

The induced combinatorial yeast-surface displayed libraries were tested separately using flow cytometry. 3 × 10^7^ cells were pelleted at 2250 rcf for 3 min and washed with 1ml of PBS+0.1%BSA at 3000 rcf for 5 min. Washed cells were incubated with antibody stains (1:25 of anti-FLAG-PE, 130-101-576, and 1:50 of anti-HA-FITC from Miltenyi, 130-120-722) for 1 hr at 4 C. Following incubation, the cells were washed with 1 ml PBS with 0.1% BSA, pelleted and then resuspended in 1 ml PBS. Samples were diluted to achieve a final concentration of 5 × 10^6^ cells/mL following which, FITC (anti-HA) and PE (anti-FLAG) intensities were detected using the Beckman Coulter Gallios flow cytometer.

Gates for cell sorting of cleaved and uncleaved populations were defined using the MoFlo Astrios Cell Sorter. Cells from the two gates – cleaved and uncleaved underwent one round of sorting and were collected until a cell count of 106 was reached. DNA was collected from each population by using a Zymoprep Kit (Omega).

## Data Availability

All related analytical results in this study are provided in the Supplementary Materials. HCV/TEV input datasets for PGCN model selection are available at Zenodo. TEV designs for PGCN screening and for flow cytometry analysis are also available at Zenodo with https://doi.org/10.5281/zenodo.7623645. All scripts to generate data and pre-trained HCV/TEV models are available in the github repository. All classification files for cleavage activities are also available in the github with https://github.com/Nucleus2014/protease-gcnn-pytorch/. All other source data are available on request from the authors.

## Code Availability

All codes and scripts to replicate PGCN results are available in https://github.com/Nucleus2014/protease-gcnn-pytorch/.

## Acknowledgments

We acknowledge NSF Molecular Foundations of Biotechnology grant CHE2226816 (to S.D.K and S.W). Joseph H. Lubin was funded by the Rutgers National Institutes of Health Biotechnology Training Program (T32 GM008339 and GM135141). Samuel Stentz was funded by the Rosetta Commons Research Experience for Undergraduates.

## Author Contributions

C.L., J.H.L., S.Z.S. and S.D.K. conceptualized the design of the study. S.D.K. and S.W. supervised the study. C.L, S.Z.S., S.W., and S.D.K. developed and designed the methodology. C.L. S.Z.S. and J.H.L. contributed to data curation. C.L. maintained research data including codes for analysis. C.L. and S.Z.S. programmed and implemented the PGCN code. J.H.L. programmed the Rosetta modelling scripts. C.L. validated models in silico and C.L. and G.W. applied statistical tools to analyze study data. V.V.S. performed yeast-based assay and flow cytometry analysis. C.L. wrote the initial draft with the help of J.H.L. and S.D.K. All co-authors contributed to writing.

## Materials & Correspondence

Correspondence to Sagar D. Khare

