## Supplementary Information for "Prediction and Design of Protease Enzyme Specificity Using a Structure-Aware Graph Convolutional Network"

Table S1A. Table for lists of cleaved and uncleaved substrates for HCV proteases (Attached)

Table S1B. Table for lists of cleaved and uncleaved substrates for TEV proteases (Attached)

Table S2. Metric Table for models based on different feature encodings for HCV and TEV data (Attached)

Table S3. Summary of Training and Test TEV Data. Constitutions of the TEV combined dataset. It shows the number of samples from cleaved or uncleaved pool split in training and test data for PGCN. Six variants with more than three mutations are simplified into 'Var', suffixed with the numbering. Var1: T146S_D148P_S153N_S170A_N177M, Var2: E107D_D127A_S135F_R203Q_K215E, Var3: T17S_N68D_E107D_D127A_F132L_S135F_F162S_K229E.

| Protease Variant | Cleaved | | | Uncleaved | | | Cleaved Percentage | Total |
|---|---|---|---|---|---|---|---|---|
| | Train | Test | Total | Train | Test | Total | | |
| WT | 286 | 116 | 402 | 85 | 31 | 116 | 78% | 518 |
| N176I | 181 | 67 | 248 | 83 | 43 | 126 | 66% | 374 |
| H28L_T30A | 30 | 10 | 40 | 29 | 9 | 38 | 51% | 78 |
| I138T_N171D_N176T | 57 | 17 | 74 | 53 | 21 | 74 | 50% | 148 |
| V209M_W211I_M218F | 98 | 38 | 136 | 64 | 26 | 90 | 60% | 226 |
| E107D_D127A_S135F | 39 | 20 | 59 | 28 | 6 | 34 | 63% | 93 |
| L2F | 785 | 339 | 1124 | 1399 | 616 | 2015 | 36% | 3,139 |
| Var1 | 124 | 64 | 188 | 73 | 26 | 99 | 66% | 287 |
| Var2 | 75 | 39 | 114 | 21 | 8 | 29 | 80% | 143 |
| Var3 | 153 | 70 | 223 | 134 | 62 | 196 | 53% | 419 |
| **Combined** | 1,828 | 780 | 2,608 | 1,969 | 848 | 2,817 | 48% | 5,425 |

Table S4. Nodes and Edges Importance Scores for HCV/TEV data (Attached)

Table S5. Table for TEV variants mutation sites indices (Attached)

Table S6. Feature descriptions for nodes and edges.

| Type | Feature | Description |
|---|---|---|
| Node | aa | One-hot encoders for amino acid types |
| | fa_intra_sol_xover | Intra-residue Lazaridis-Karplus solvation energy |
| | fa_intra_rep | Intra-residue Lennard-Jones repulsive energy |
| | rama_prepro | Ramachandran preference score of backbone angles |
| | omega | Omega dihedral score |
| | p_aa_pp | Probability of amino acid type at backbone angles |
| | fa_dun | Side-chain conformation score |
| | ref | Reference potential |
| | is_substrate | 1 if the node belongs to the substrate; 0 otherwise |
| Edge | fa_atr | Lennard-Jones attractive potential |
| | fa_rep | Lennard-Jones repulsive potential |
| | fa_sol | Lazaridis-Karplus solvation potential |
| | lk_ball_wtd | Asymmetric solvation potential |
| | fa_elec | Coulombic electrostatic potential |
| | hbond | Hydrogen bonding potential |
| | intermolecular | 1 if the edge is between a substrate residue and a protease residue; 0 otherwise |
| | covalent_bond | 1 if pairwise residues form a covalent bond; 0 otherwise |

Table S7A. Table for graph indices for HCV proteases

| Node Index | PDB index | Substrate | AA Type (three-code) | AA Type (one-code) |
|---|---|---|---|---|
| 1 | 198 | TRUE | | |
| 2 | 199 | TRUE | | |
| 3 | 200 | TRUE | | |
| 4 | 201 | TRUE | | |
| 5 | 202 | TRUE | | |
| 6 | 58 | FALSE | PHE | F |
| 7 | 70 | FALSE | VAL | V |
| 8 | 72 | FALSE | HIS | H |
| 9 | 73 | FALSE | GLY | G |
| 10 | 96 | FALSE | ASP | D |
| 11 | 138 | FALSE | ARG | R |
| 12 | 147 | FALSE | ILE | I |
| 13 | 150 | FALSE | LEU | L |
| 14 | 151 | FALSE | LYS | K |
| 15 | 152 | FALSE | GLY | G |
| 16 | 154 | FALSE | SER | S |
| 17 | 170 | FALSE | ARG | R |
| 18 | 171 | FALSE | ALA | A |
| 19 | 172 | FALSE | ALA | A |
| 20 | 173 | FALSE | VAL | V |
| 21 | 174 | FALSE | CYS | C |
| 22 | 175 | FALSE | THR | T |
| 23 | 176 | FALSE | ARG | R |
| 24 | 177 | FALSE | GLY | G |
| 25 | 178 | FALSE | VAL | V |
| 26 | 179 | FALSE | ALA | A |
| 27 | 180 | FALSE | LYS | K |
| 28 | 181 | FALSE | ALA | A |
| 29 | 182 | FALSE | VAL | V |
| 30 | 183 | FALSE | ASP | D |
| 31 | 197 | TRUE | | |
| 32 | 203 | TRUE | | |
| 33 | 204 | TRUE | | |
| 34 | 205 | TRUE | | |

Table S7B. Table for graph indices for TEV proteases

| Node Index | PDB index | Substrate | AA Type (three-code) | AA Type (one-code) |
|---|---|---|---|---|
| 1 | 302 | TRUE | | |
| 2 | 303 | TRUE | | |
| 3 | 304 | TRUE | | |
| 4 | 305 | TRUE | | |
| 5 | 306 | TRUE | | |
| 6 | 30 | FALSE | | T |
| 7 | 32 | FALSE | | L |
| 8 | 44 | FALSE | | N |
| 9 | 45 | FALSE | | K |
| 10 | 46 | FALSE | | H |
| 11 | 47 | FALSE | | L |
| 12 | 49 | FALSE | | R |
| 13 | 81 | FALSE | | D |
| 14 | 134 | FALSE | | S |
| 15 | 139 | FALSE | | F |
| 16 | 148 | FALSE | | D |
| 17 | 149 | FALSE | | G |
| 18 | 151 | FALSE | | C |
| 19 | 152 | FALSE | | G |
| 20 | 167 | FALSE | | H |
| 21 | 168 | FALSE | | S |
| 22 | 169 | FALSE | | A |
| 23 | 170 | FALSE | | S |
| 24 | 171 | FALSE | | N |
| 25 | 172 | FALSE | | F |
| 26 | 173 | FALSE | | T |
| 27 | 174 | FALSE | | N |
| 28 | 175 | FALSE | | T |
| 29 | 176 | FALSE | | N |
| 30 | 177 | FALSE | | N |
| 31 | 178 | FALSE | | Y |
| 32 | 208 | FALSE | | S |
| 33 | 209 | FALSE | | V |
| 34 | 211 | FALSE | | W |
| 35 | 213 | FALSE | | G |

| | | | | |
|---|---|---|---|---|
| 36 | 214 | FALSE | | H |
| 37 | 215 | FALSE | | K |
| 38 | 216 | FALSE | | V |
| 39 | 217 | FALSE | | F |
| 40 | 218 | FALSE | | M |
| 41 | 219 | FALSE | | S |
| 42 | 220 | FALSE | | K |
| 43 | 221 | FALSE | | P |
| 44 | 301 | TRUE | | |
| 45 | 307 | TRUE | | |
| 46 | 308 | TRUE | | |
| 47 | 309 | TRUE | | |

Table S8. Yeast-assay experimental and PGCN predicted probabilities for TEV designs (Attached)

Table S9. Table for the percentage of substrate overlaps

| Overlap Percentage | | Test set for | | | |
|---|---|---|---|---|---|
| | | WT | A171T | D183A | Triple |
| Training set for | WT | 0.00% | 2.42% | 1.53% | 1.09% |
| | A171T | 1.32% | 0.00% | 2.88% | 2.02% |
| | D183A | 1.01% | 2.92% | 0.00% | 1.79% |
| | Triple | 1.21% | 4.11% | 2.91% | 0.00% |

Table S10. Cross Test Table for PGCN models based on HCV data (Attached)

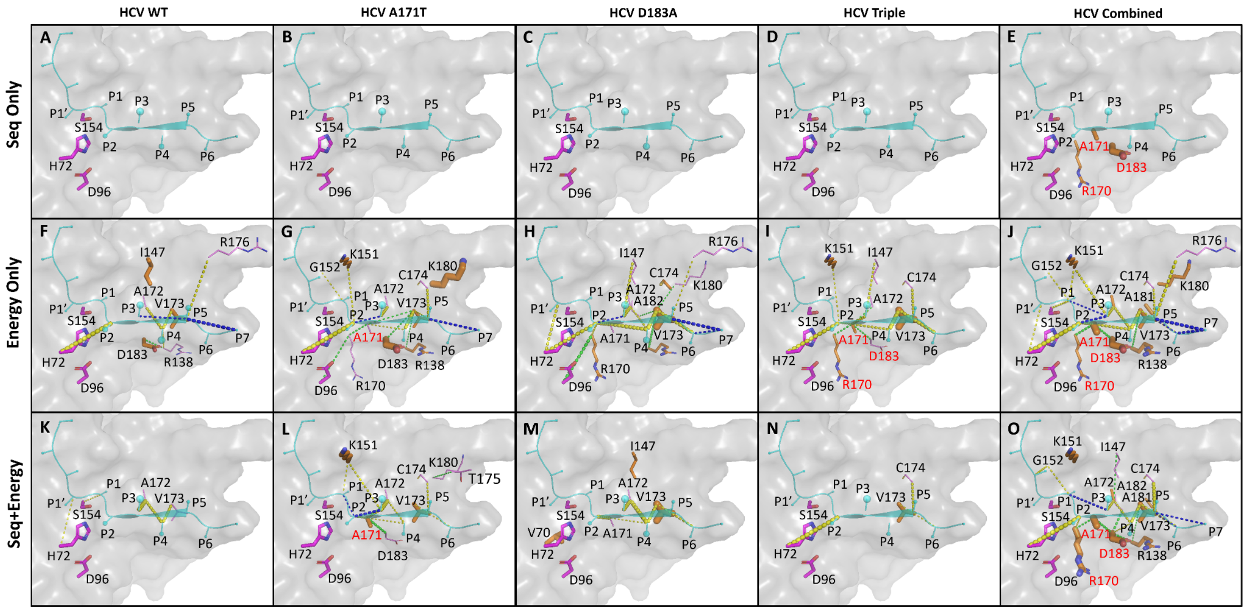


Figure S1. Structural depictions of node and edge importances in cleavage classification for HCV data.

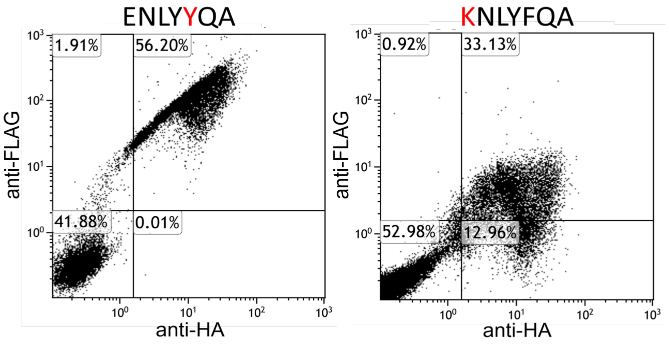


Figure S2. 2-D scatter plots (anti-FLAG on y-axis; anti-HA on x-axis) for wild-type TEV on Y at P2 (left) or K at P6 (right). The corresponding FLAG/HA ratios for these plots are >0.4 which is our cutoff for cleaved constructs.

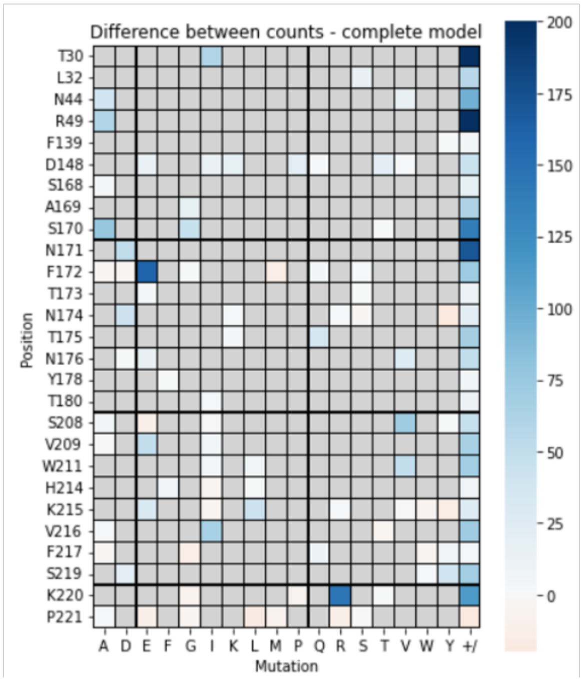


Figure S3. Heatmap for Single mutations for raw TEV protease designs.

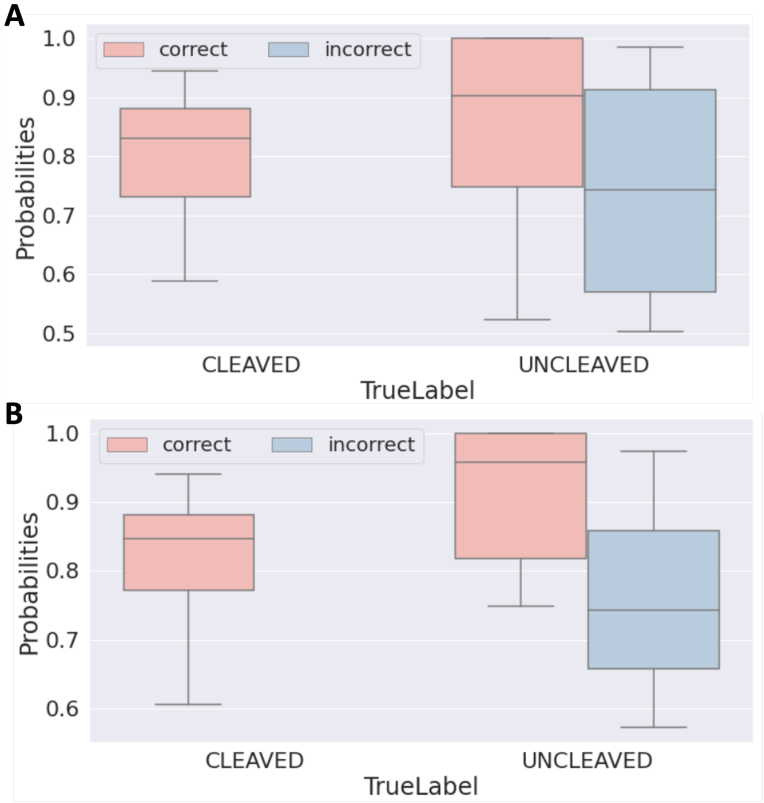


Figure S4. Robustness of predictions among decoys. A. Ten decoys per design; B. Best decoys for designs

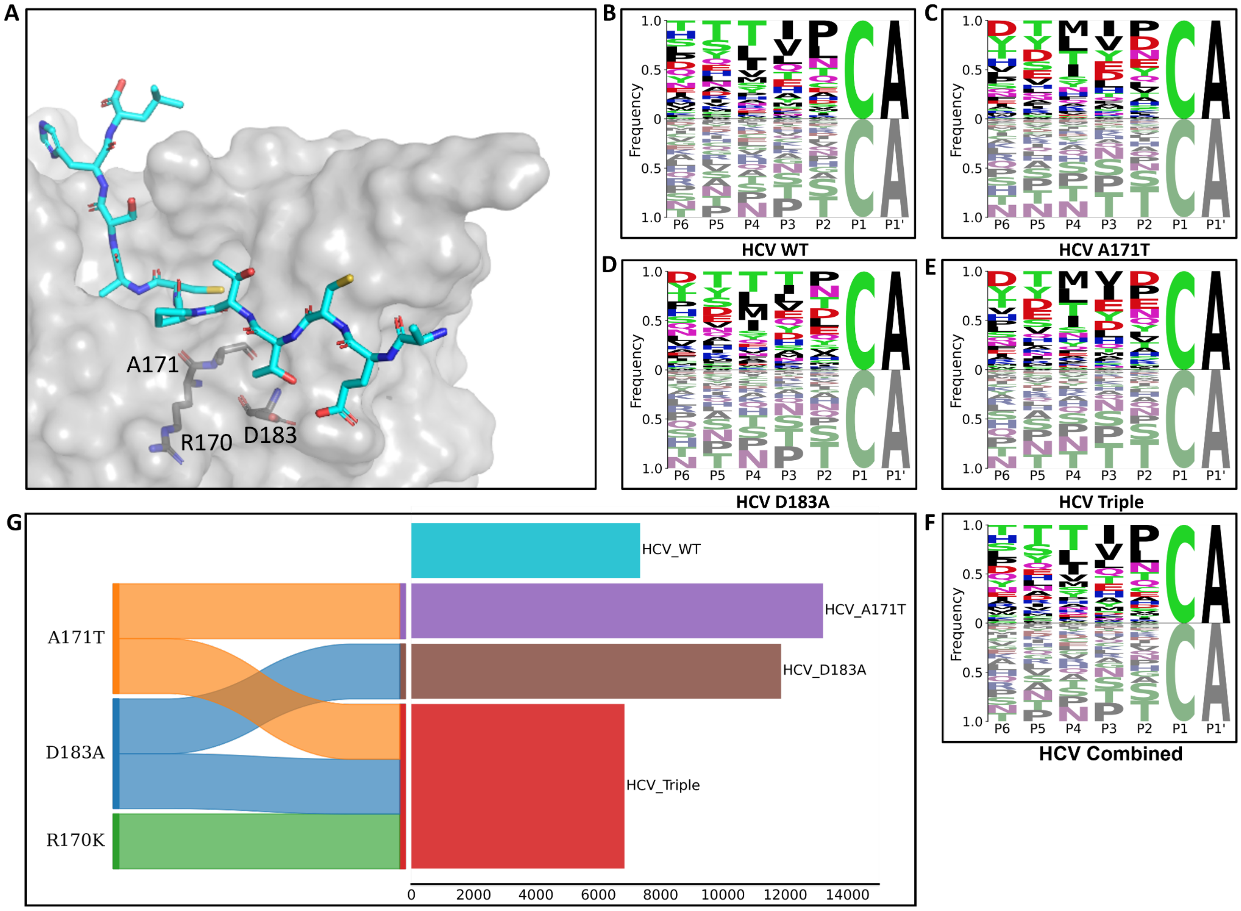


Figure S5. HCV input data variety of mutations and sequence. A) Mutation sites (grey) in HCV-PR protease structure are around the substrate (cyan); B-F) Sequence logo plots of input data for WT, A171T, D183A and Triple (mutation sites: A171T, D183A, R170K). Logo above x-axis shows frequencies of each amino acid type per position for cleaved sequences, while logo before x-axis shows frequencies of each amino acid type per position for uncleaved sequences; G) Sankey diagram with the barplot, showing variability for each HCV variant input data. Sankey diagram shows mutations (on the left) of each HCV variant (on the right), while the barplot on the right shows the number of substrates for each variant.

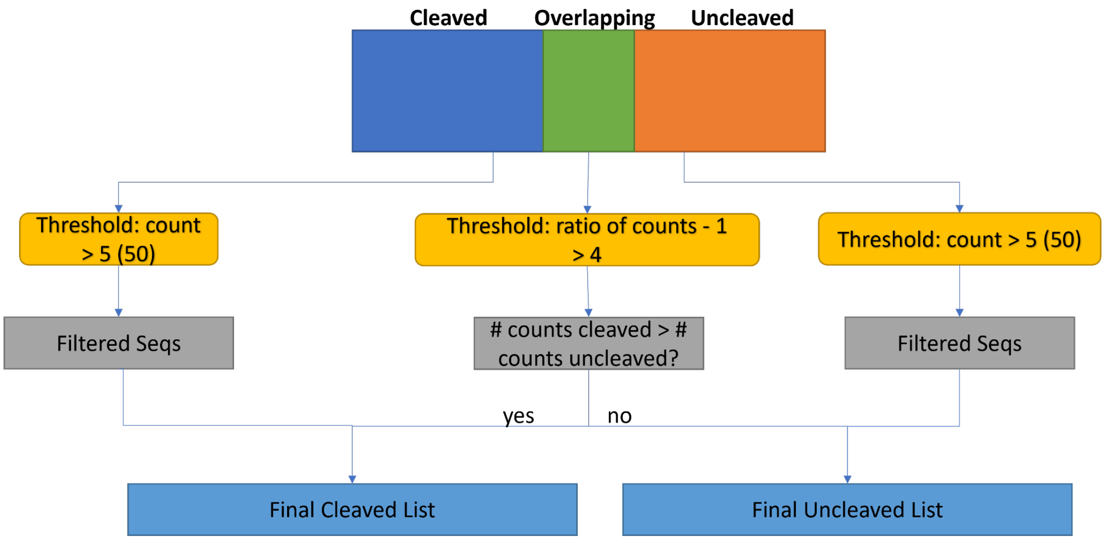


Figure S6. The pipeline of preprocessing deep sequencing TEV data and filtering out cleavage lists for the PGCN model.

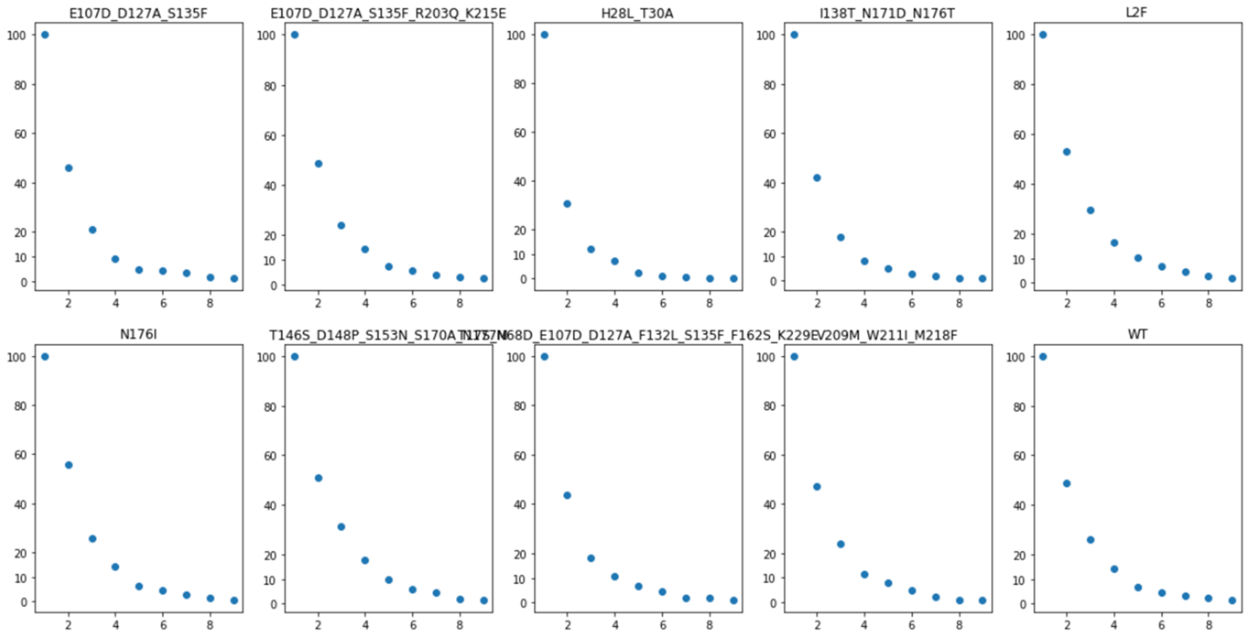


Figure S7. The threshold selection for the overlapping samples between deep cleaved and uncleaved single mutated libraries.

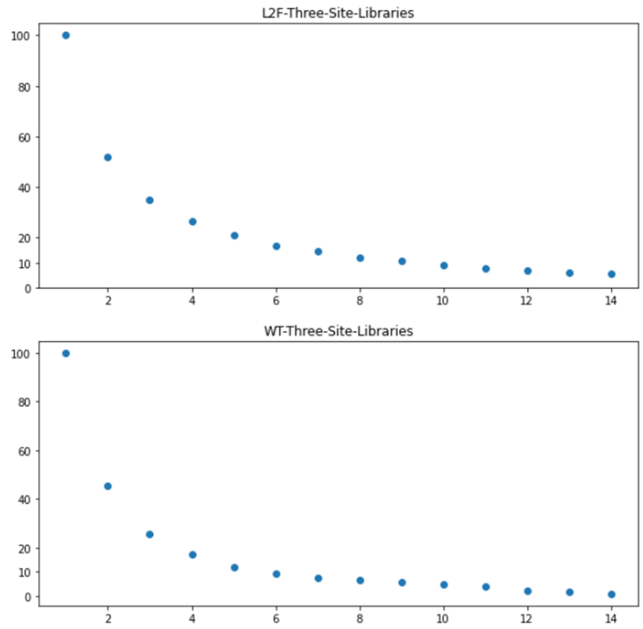


Figure S8. The threshold selection plot for the overlap between deep cleaved and uncleaved triple mutated libraries.

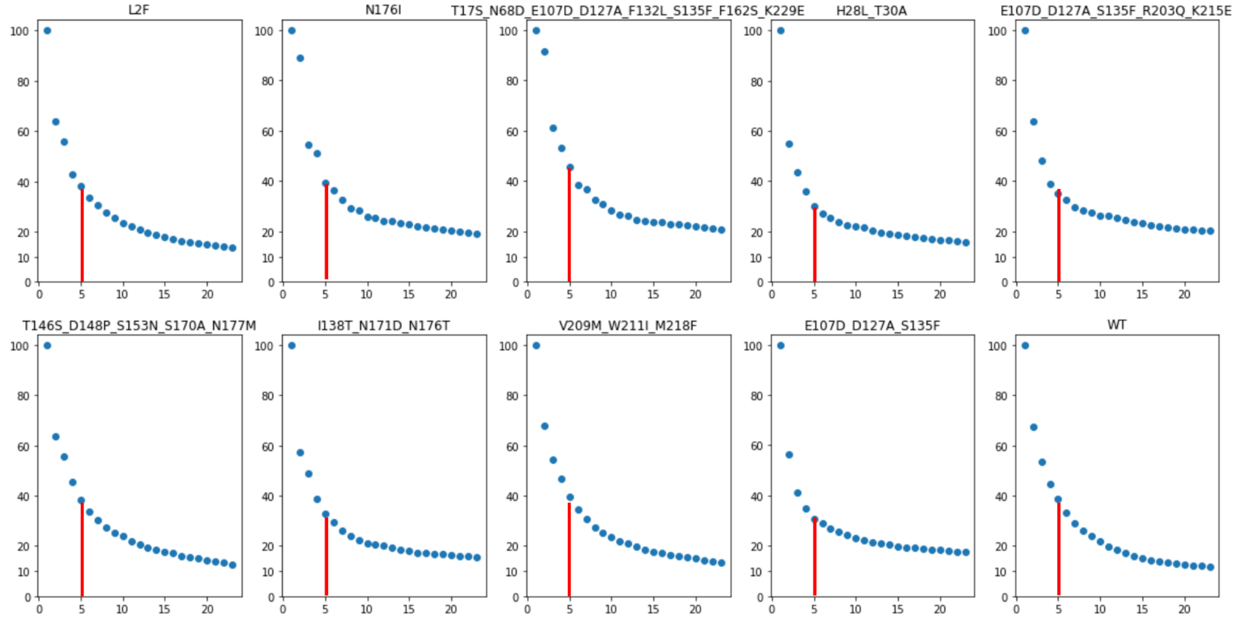


Figure S9. The threshold selection for the non-overlap between deep cleaved and uncleaved single mutated libraries.

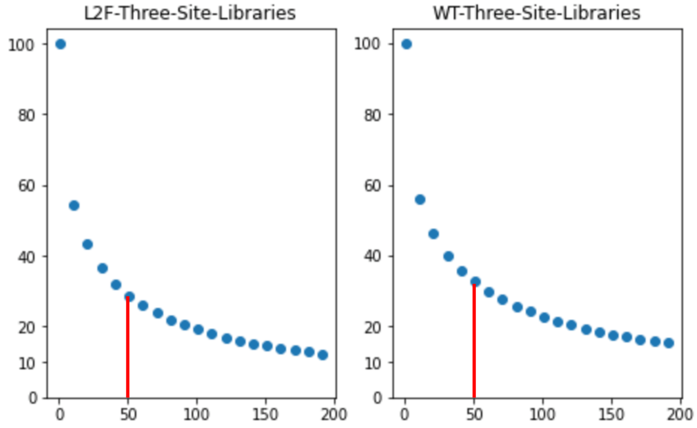


Figure S10. The threshold selection for the non-overlap between deep cleaved and uncleaved triple mutated libraries.

### Supplementary Methods

#### *HCV Experimental Data Preparation*

The HCV experimental data used to train the PGCN were from a previously published study[32] in which a large library of candidate substrates with variation in the P6-P2 region was screened for cleavage by WT HCV and three drug-escape mutants, with substitutions A171T, D183A, and R170K+A171T+D183A respectively. Experimental classification of cleavage was performed by yeast surface display[44], with each substrate assigned as either cleaved or uncleaved in binary classification.

The HCV dataset was generated using yeast surface display with a construct in which a P6-P4' substrate was expressed between two antibody binding sites, resulting in the loss of the second antibody signal for cleaved substrates. Yeast cells expressing cleaved substrates were separated from those expressing uncleaved substrates by fluorescence-activated cell sorting (FACS), and the separated pools were deep sequenced to determine all substrates within the pool.

#### *TEV Deep Sequencing Data Processing*

Substrate specificities from deep sequencing are an excellent resource for preparing input substrate lists for one or multiple proteases ready for PGCN model training. However, before sending sequences to deep sequencing, experiments often introduce errors in identifying cleavage activity. Therefore, we developed a pipeline to preprocess deep sequencing data from phage substrate display experiments collected from Packer et al., 2017 and filter sequences with promising cleavage information. The pipeline is applicable for analyzing deep sequencing data from cleavage activity determination experiments. First, substrate counts in all libraries from each wild type or variant are normalized by a normalized factor of the number of sequences in the most extensive library divided by the number of sequences in the target file. Therefore, normalized counts in different libraries are comparable and can be summed up for further processing. Second, for each protease type, libraries are combined into two large populations, based on cleaved or uncleaved sequences, where normalized counts of redundant sequences across different libraries are summed up, and new sequences are added directly with their specific counts in the corresponding library.

We assume little overlap between cleaved and uncleaved sequence populations, so we speculate the optimal cutting threshold of normalized counts by analyzing the trending of the overlap between two populations scaled to initial overlap (when the threshold is set to 1) as the threshold increases. Based on the assumption, we process data separately into three sub-parts (**Figure S6**), the overlapping region that contains sequences both in cleaved and uncleaved populations, and two non-overlapping regions that contain sequences only in cleaved or uncleaved populations.

The definition of the threshold for overlapping sequences is the ratio of normalized counts of cleaved sequences and uncleaved sequences minus 1. The optimal threshold is set to 4 for data of single mutation sites on substrates, meaning sequences can be grouped as cleaved if their normalized counts in the cleaved pool are 5 times greater than those in the uncleaved pool (**Figure S7**). For three mutation sites on substrates, 6 is set for WT data, and 10 is for L2F data (**Figure S8**).

The remained non-overlapping data in both cleaved and uncleaved pools are filtered based on a threshold of normalized counts directly, which are 5 for variant data of single mutation sites on substrates, and 50 for variant data of three mutation sites on substrates. Those two optimal thresholds ensure a sudden decrease of overlaps between cleaved and uncleaved libraries for all TEV protease types (**Figure S9 & S10**) and enough data points within the threshold to be fed into PGCN.

##### *Computational design process for TEV*

Binding pocket residues (K141, N171, T173, T175, N176) interacting with P6 or those residues interacting with P2 i.e. V209, W211 and M218 were computationally redesigned using Rosetta FastDesign to favor specificity toward K at the P6 position and Y at P2, respectively.

TEV redesign was performed in three phases. The first phase aimed to identify potential affinity-enhancing substitutions with high throughput. It followed a similar protocol to protease-substrate complex model generation, though the amino acid exchange was allowed for the substrate-interfacing residues, and minimization was applied to a second shell of protease residues that interface with the designable set. The sequences (p6: KNLYFQ/A, p2: ENLYYQ/A) full substrate were used as design targets. 1,000 design decoys were generated. In the second phase, generated models were manually reviewed to identify promising substitutions for experimental validation, and several rational substitutions were also added to the pool, which results in 280 designs for p2 and 4,320 designs for p6. Energetically favorable designs of 96 targeting P6 and 18 targeting P2 are then selected manually, while PGCN selected a less stringent set, with 126 of 4,320 P6-targeted and 200 of 280 P2-targeted designs.

The third phase included the creation of a combinatorial library of all selected substitutions at all variable positions. This library was generated both experimentally and computationally. In the case of the computational library generation, we used the same protease-substrate complex model generation protocol, with the addition of making any protease substitutions prior to FastRelax minimization.

##### *Exploring PGCN Generalizability by Cross-Test Analysis*

To design rational protease designs is based on their specificity towards specific substrates. Therefore, having a tool to quickly reveal protease specificity is very helpful for the efficiency of protease designs. In the previous section, all PGCN models trained on one of the HCV datasets are tested on the same dataset. It is unknown how PGCN performs on new mutations unseen by the training datasets. Herein, To investigate if the models learned by PGCN for a single variant protease could be further generalized to protease variants outside the training data, we cross-tested prediction accuracy for each PGCN model trained on one of five HCV datasets on the other four. Each PGCN model used for the cross-test analysis is selected from three models with the same parameters but different random seeds based on their accuracy being the median of the three models. To avoid data leaking, we removed the testing samples that overlap substrates with the training dataset, even though the total number of overlaps is very small (**Table S9**). In **Table S10**, the PGCN model trained on the combined set always had lower accuracy than the model trained on the single variant set (such as 90.57% vs. 96.37% tested on A171T). Therefore, seeing more mutations does not guarantee significantly better performance.

However, if we only focus on cross-test tasks, it turns out that the PGCN model still benefits from observing the specific mutation to predict specificity rationally, since models trained on the combined set perform the best measured by different metrics for most cross-test tasks. For example, when we consider models using sequence and energy features, for the cross-test on A171T, the model trained on the combined set reached 90.57% accuracy, and 82.76% F1 score, much higher than other models except the one trained on A171T itself.

However, the PGCN model trained on a single protease still has a certain ability to infer a limited variety of protease specificity. Whatever feature encoding is used, the model trained on A171T performed the prediction on the Triple mutant (such as 87.69% accuracy, 84.24% F1, 83.82% AP, 94.36% AUC using sequence+energy features) comparable to the best baseline model trained on Triple (SVM: 91.96% accuracy, 91.45% F1, 89.06%AP, 96.22% AUC) (**Table S2**).
